# Accounting for fast vs slow exchange in single molecule FRET experiments reveals hidden conformational states

**DOI:** 10.1101/2024.06.03.597137

**Authors:** Justin J. Miller, Upasana L. Mallimadugula, Maxwell I. Zimmerman, Melissa D. Stuchell-Brereton, Andrea Soranno, Gregory R. Bowman

## Abstract

Proteins are dynamic systems whose structural preferences determine their function. Unfortunately, building atomically detailed models of protein structural ensembles remains challenging, limiting our understanding of the relationships between sequence, structure, and function. Combining single molecule Förster resonance energy transfer (smFRET) experiments with molecular dynamics simulations could provide experimentally grounded, all-atom models of a protein’s structural ensemble. However, agreement between the two techniques is often insufficient to achieve this goal. Here, we explore whether accounting for important experimental details like averaging across structures sampled during a given smFRET measurement is responsible for this apparent discrepancy. We present an approach to account for this time-averaging by leveraging the kinetic information available from Markov state models of a protein’s dynamics. This allows us to accurately assess which timescales are averaged during an experiment. We find this approach significantly improves agreement between simulations and experiments in proteins with varying degrees of dynamics, including the well-ordered protein T4 lysozyme, the partially disordered protein apolipoprotein E (ApoE), and a disordered amyloid protein (Aβ40). We find evidence for hidden states that are not apparent in smFRET experiments because of time averaging with other structures, akin to states in fast exchange in NMR, and evaluate different force fields. Finally, we show how remaining discrepancies between computations and experiments can be used to guide additional simulations and build structural models for states that were previously unaccounted for. We expect our approach will enable combining simulations and experiments to understand the link between sequence, structure, and function in many settings.

## Introduction

A protein’s function is determined by the ensemble of structures that it adopts^1–6^, but building atomically-detailed models of these ensembles to probe ensemble-function relationships remains challenging^7^. Of course, the high-resolution structures that structural biologists have become adept at solving are of enormous value. Despite this, ongoing challenges with tasks like drug and protein design highlight the limits of the structure-function paradigm ^8^. We expect having detailed models of the rest of a protein’s structural ensemble would lead to dramatic improvements in our understanding and ability to design such systems^9–11^. Unfortunately, most of the alternative structures a protein can adopt are difficult to detect and/or characterize experimentally because they have too low of a probability (i.e. high energy)^12–14^.

Single molecule Förster resonance energy transfer (smFRET) experiments are a powerful tool for studying the distribution of structures that a protein adopts, including high energy states that are invisible to many other techniques ^15–17^. In these experiments, a donor and an acceptor fluorophore are attached to two different residues in a protein. The donor fluorophore on a single protein is then excited and one measures how many acceptor and donor photons are emitted ^18,19^. The probability of transferring energy from the donor to the acceptor fluorophore, called the FRET efficiency, reports on a variety of valuable structural properties, including the distance between the fluorophores, their relative orientations, and the timescale on which they are rotating. Making many measurements results in a probability distribution of FRET efficiencies. These FRET efficiency distributions report on the distribution of structures the protein adopts and have proved to be a powerful means of revealing the conformational heterogeneity of proteins ^20–23^.

Unfortunately, one cannot extract atomically-detailed structural models from smFRET data in a manner analogous to fitting structures to electron density from crystallography or cryoEM. smFRET data is inherently sparse, with each experiment reporting on the structure and dynamics of a single pair of dyes. One can perform experiments for multiple dye positions to learn about more of the protein structure. However, each experiment is independent, making it hard to discern any correlations between the behavior of different parts of the protein. While multi-color FRET experiments are being developed^24^, they are quite challenging to perform and still can’t measure many distances in parallel. Another challenge is that there is not a one-to-one mapping between the FRET efficiency and the distance between a pair of residues.

Combining atomically detailed computer simulations with smFRET experiments could yield experimentally grounded models of protein conformational ensembles with the desired resolution^25,26^. Ideally, there would be a method to predict energy transfer distributions from simulations to show that these predictions were in perfect agreement with smFRET experiments. Then one could analyze the simulations, making use of the atomistic structural and dynamical information they provide to generate new hypotheses, and test those hypotheses experimentally.

While there are cases where smFRET experiments and simulations are in good agreement, the agreement between the two approaches is often limited ^25^. A variety of approaches have been employed to close this gap. For example, scaling factors have been used to shift computational predictions into closer alignment with experiments ^27,28^. Others have employed reweighting schemes to shift the relative probabilities of structures from their simulations and bring their predictions into closer agreement with experiments ^29–31^. There have also been efforts to develop improved methods for predicting the probability of energy transfer from simulations (e.g. by modeling in the dyes) and to improve force fields ^26,28,32–42^. However, there is still room for improvement.

Here, we explore the importance of accounting for kinetic effects in smFRET experiments when connecting with simulations. Each FRET efficiency measured in an smFRET experiment is the ratio of acceptor photons to all photons emitted during some time interval. This time interval typically ranges from one to ten milliseconds depending on the experimental setup (e.g. TIRF vs diffusion confocal, laser power, etc.) ^22,23^. It has long been recognized in the smFRET community that this means each FRET efficiency measured is, therefore, averaging across whatever conformational dynamics occur during the one to ten millisecond time interval. As in NMR experiments, conformations that are exchanging more quickly than this measurement time will be averaged together, while conformations that are exchanging more slowly will not. Significant effort has gone into dealing with this time-averaging when analyzing experiments ^43–53^. For example, it is common to perform global fits to many measurements with different dye positions and solvent conditions ^54–56^, fit hidden Markov models to photon traces ^21,46,51^, or dissect the correlation between FRET efficiency and other fluorescence observables that report on shorter timescales ^44,45,55^. New experimental approaches are also being developed to shorten the timescale over which FRET efficiencies are measured ^57–61^. Nonetheless, accounting for time-averaging could dramatically improve agreement between experiments and simulations enabling these two approaches to be used even more effectively to advance our understanding of the ensemble-function relationship.

We present an approach for accounting for time-averaging when predicting FRET efficiencies from simulations and assess its performance on three well-studied systems that exemplify different extents of dynamics. We start with Apolipoprotein E4 (ApoE4), as it contains both ordered and disordered regions and addresses the applicability of our approach to each ^54,62^. We also apply our approach to T4 Lysozyme, a well-ordered system that has recently been extensively characterized using 33 distinct smFRET labeling positions ^55^.

Finally, we apply our approach to amyloid-β40 (Aβ40), a 40 amino acid highly disordered protein ^28^. One recent study produced simulations of Aβ40 with a variety of force fields, giving us the chance to test how well different force fields perform when combined with our approach for accounting for time-averaging when predicting the experimentally observed energy transfer distribution ^63^.

## Results

### Accounting for time-averaging dramatically improves agreement between simulations and experiments for a partially-disordered protein

We developed an approach for predicting FRET efficiencies from simulations in a manner that accounts for time-averaging by drawing on Markov state models (MSMs) ^64–67^ built from molecular dynamics simulations. An MSM is a network model that describes a molecule’s conformational space in terms of the structural states it adopts and the probabilities of hopping between every pair of states in a fixed time interval. These models integrate information from many independent simulations to capture length and time scales that are far beyond the reach of any individual simulation. Importantly, we can use an MSM to generate a synthetic trajectory using a kinetic Monte Carlo scheme in which one chooses a random starting state and then iteratively adds new random states based on the transition probabilities from the current state to all other states. To mimic a smFRET experiment, we use one MSM to describe the protein’s conformational dynamics and separate MSMs for each of the dyes. First, we select a random experimental photon time trace and use our protein MSM to generate a synthetic trajectory of the same length. Then we identify conformations in our synthetic trajectory that correspond to the times photons were detected in the experiment. We assume these are the conformations that emit photons and then choose whether to label each photon as coming from the donor or acceptor dye as follows. First, we generate a set of plausible dye conformations by mapping representative structures from each state in our dye MSM onto the protein structure and removing any that form steric clashes with the protein.

Next, we use our dye MSMs to simulate the dynamics of the dye (on a fixed protein structure) leading up to emission of a photon. At each step of these dye simulations, we use a Monte Carlo move to decide if the donor emits a photon, transfers energy leading to emission of an acceptor photon, or stays excited similar to previous efforts ^40,41^. If the dye remains excited, both dyes are allowed to hop to another state in the MSM. We repeat this process for the remainder of the synthetic trajectory to simulate a photon burst, returning the average FRET efficiency, or the number of acceptor photons divided by the total photons, for that burst.

Finally, we repeat this process over multiple trajectories until an adequate number of photon bursts have been sampled. Since we model dyes as a post-processing step instead of including the dyes in the simulations, it is easy to scale this approach to predict the observed FRET for many dye positions. Furthermore, simulating the dye dynamics allows us to minimize the number of adjustable parameters, as we do not need to select constant values like a Förster radius that are required by other approaches.

To test our approach, we applied it to the partially disordered protein ApoE4. Our recent work presented smFRET measurements for five different pairs of dye positions on this protein. Some of these dye positions report on dynamics within the largely folded N-terminal domain while others report on the partially disordered and highly flexible C-terminal domain^54^. Therefore, comparisons between simulations of this protein and experiments speak to the utility of our approach for both well-folded and disordered structures. We previously showed that ApoE4 predominately adopts three conformational states: a closed state, an open state, and an extended state. Identifying these three states experimentally required an enormous number of measurements, including multiple dye positions each at varying levels of denaturant. Even with this wealth of data, building atomically detailed structural models of the different states required extensive molecular dynamics simulations, totaling over 3 ms of aggregate simulation, which we showed were in reasonable agreement with experiments using a simplified version of the approach presented here. That approach did not model dye dynamics and, therefore, required us to choose constant values for parameters like the fluorescence anisotropy used in the Förster radius.

Assessing different ways of predicting FRET efficiencies from simulations and examining the actual distance distributions in those simulations highlights the importance of accounting for time-averaging (Figure 1). For example, Figure 1A shows the modeled inter-dye distance distribution between residues 5 and 86 in the folded, N-terminal domain. This distribution has two peaks, which roughly correspond to the closed and open states of ApoE4 (Figure 1B). If one assumes that smFRET measurements are instantaneous (i.e. there is no time-averaging), then the distribution of FRET efficiencies that one predicts retains these two peaks. However, accounting for time-averaging causes these two peaks to collapse into a single peak because the different populations are in fast exchange (Figure 1C). Importantly, the FRET efficiency we predict by accounting for time-averaging is in good agreement with the experimental data. Without accounting for time-averaging, we would have come to the erroneous conclusion that our simulations were in poor agreement with experiments. By accounting for time-averaging, we instead find good agreement with experiments and can use the simulation data to help identify the different populations that give rise to the experimentally observed smFRET data.

**Figure 1:**
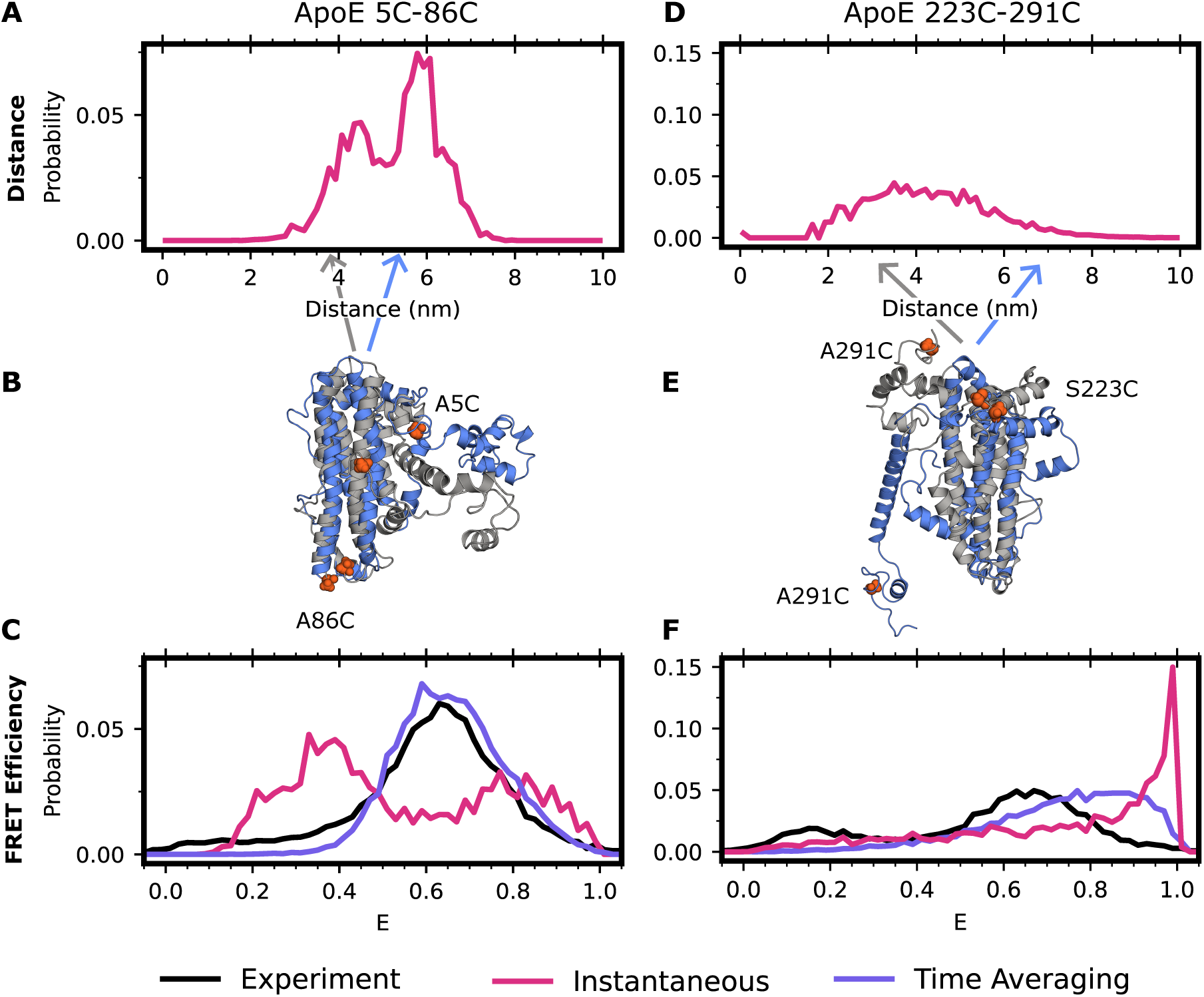
Accounting for time averaging significantly alters the apparent structural distribution from our model and increases agreement with experiments. (A) Inter-dye distances for apolipoprotein E labeled with Alexafluor 488 and Alexafluor 594 at positions 5 and 86 or (D) 223 and 291. In red is the equilibrium (instantaneous) distribution accounting for the distance added or subtracted by dye positioning (B,E) Exemplar structures of ApoE at two distinct dye-distance positions. Arrows indicate the portion of the distance distribution the structure occupies. (C) FRET efficiencies obtained for positions 5 and 86 or (F) 223 and 291. In black is the experimental distribution, in red is the result when not accounting for conformational dynamics of ApoE, and in purple is the time-averaged trace of the red trace.

Repeating the analysis above for the other dye positions supports the importance of accounting for time-averaging. For example, the modeled inter-dye distance distribution between residues 223 and 291 in the disordered C-terminal domain is broad and symmetrical (Figure 1D). The distribution of FRET efficiencies one would predict without accounting for time-averaging is skewed to large FRET efficiencies, in poor agreement with the experimentally observed distribution of FRET efficiencies. Accounting for time-averaging improves the agreement between simulations and experiments (Figure 1F). Importantly, the MSM accounts for motions occurring over multiple timescales enabling us to automatically average together states which are interconverting rapidly in a single energy basin while simultaneously capturing the differences between states that are interconverting slowly and thus broadening the histogram. Similar results are found for other dye positions (Figure S1). Treating the dyes as a point cloud rather than modeling their dynamics also gives similar results (Figure S2), though this approach requires the choice of a constant Förster radius that can be a source of error if a poor choice is made or if the dyes are not isotropically rotating.

### Time-averaging improves experiment-simulation agreement for the entire spectrum from ordered to disordered proteins

Given that ApoE has a mix of ordered and disordered regions, we reasoned that our time-averaging approach should be equally applicable to fully ordered and disordered systems. To test this hypothesis further, we used our approach on the highly ordered protein, T4 lysozyme, and the intrinsically disordered protein (IDP), Aβ40. Both T4 lysozyme and Aβ40 benefit from a plethora of prior structural studies, including experimental smFRET characterization ^28,55^. For Aβ40, we make use of an existing 30 μs long simulation in the amber99sb forcefield, which was found to match NMR order parameters reasonably well ^63^. For lysozyme, we performed 5 independent 5 μs long all-atom molecular dynamics simulations in explicit tip3p solvent and the amber03 force field as described in the methods section. For both Aβ40 and lysozyme, we clustered our datasets, made MSMs, modeled on the appropriate dye pairs to match the experimental setup, and investigated the predicted FRET efficiencies using our time-averaging approach.

As expected, we found that accounting for time-averaging is important for both systems (Figure 2). We first calculate the lysozyme FRET efficiency for one of the experimental FRET probe distances, residues 44-150, using Alexa 488 maleimide and Alexa 647 hydroxylamine dyes. We find strong agreement between our time-averaging approach and the experimental results for lysozyme (Figure 2A). We note that both time-averaging and instantaneous FRET are in reasonable agreement with experimental data for this probe position, though both miss a population at high FRET efficiency. We next calculate the FRET efficiency for Aβ40 using Alexa 488 hydroxylamine and Alexa 647 maleimide attached to positions 1 and 40. We find that the results significantly improve upon the distribution obtained without time-averaging. However, the distribution is shifted overall towards higher FRET efficiencies, suggesting either insufficient sampling or force fields issues (Figure 2B). Overall, these findings demonstrate that accounting for time-averaging is helpful when there are conformations in fast exchange and is equivalent to other approaches when such exchange is absent.

**Figure 2:**
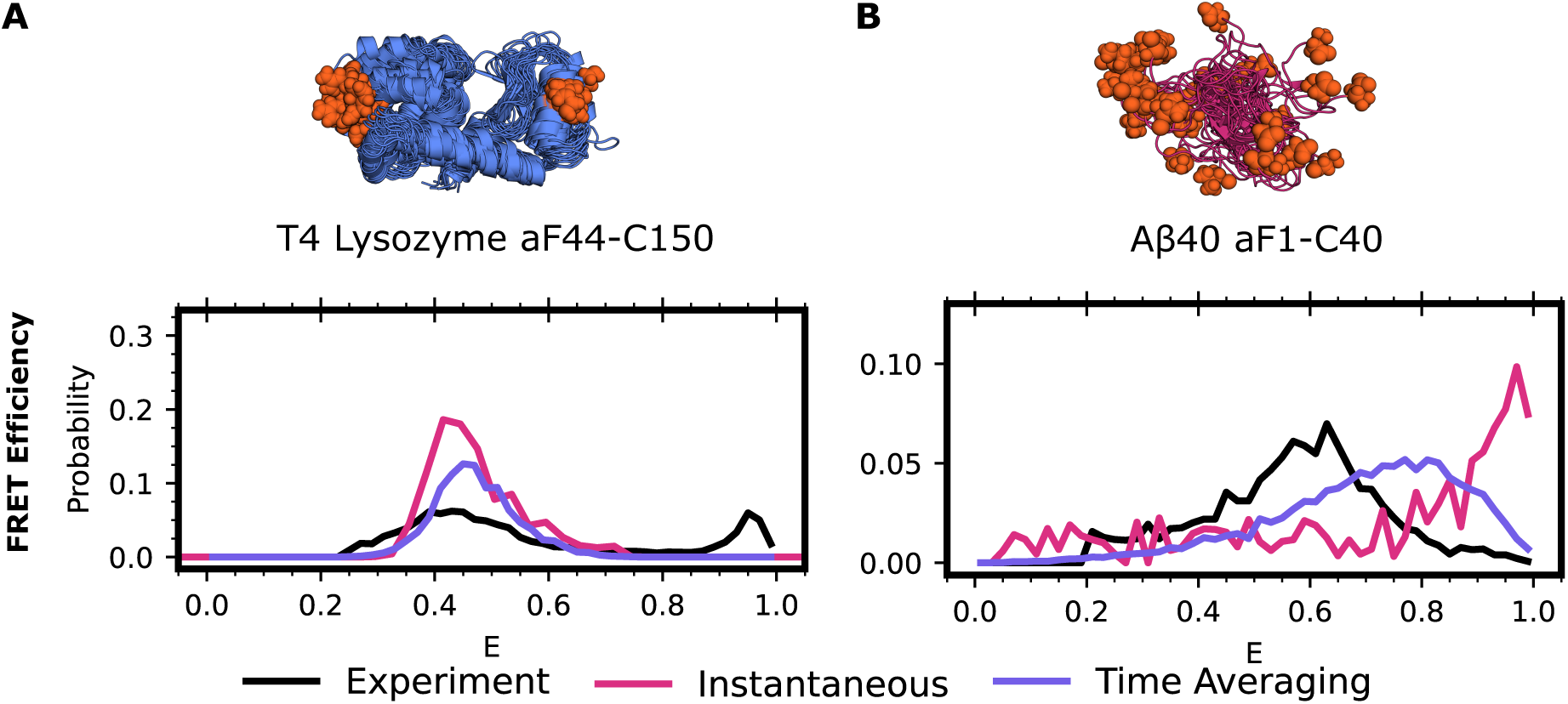
smFRET time averaging impacts proteins across the ordered. A) FRET efficiencies for T4 Lysozyme labeled at 44 (*para*-acetylphenylalanine) and 150 (cysteine) with Alexa 488 and Alexa 647 or B) Aβ40 labeled at positions 1 (*para*-acetylphenylalanine) and 40 (cysteine) with Alexa 488 and Alexa 647. In black is the experimental distribution, red the result when not accounting for protein dynamics, and purple accounting for protein dynamics via time-averaging. Protein structures are the 15 most probable states in the MSM with labeling positions indicated in orange spheres. Experimental donor only counts (E < 0.25) have been removed for ease of comparison.

### Directing sampling based on discrepancies between predicted and observed FRET reveals a novel conformation of lysozyme

Given the strong agreement between our predicted energy transfer distributions and experiments for folded and partially disordered systems, we reasoned that remaining discrepancies may point to under sampled regions of conformational space in simulations. Indeed, the prior study on lysozyme concluded that the minor population could not be explained by any structure of lysozyme existing in the protein data bank ^55^. If this is true, then we should be able to improve the agreement between simulations and experiments by driving simulations to sample structures with FRET values that are not observed often enough compared to experiments.

To explore this possibility, we sought to provide a structural model for a minor population that was previously observed in an extensive smFRET study of lysozyme. That study presented smFRET measurements for 33 pairs of dye positions. For 17 of these dye pairs, the authors observe a minor state in the FRET efficiency that they could not explain based on any of the numerous published crystal structures of this protein. When probing residues 44-150, this minor population has high FRET efficiency, a metric which would require the dyes to come closer together than is conceivable based on a clamshell motion of the two lobes of lysozyme. Furthermore, we would not expect our simulations (aggregate simulation time of 25 μs) to reach this minor state since the experiments suggest that it is accessed with a rate of ∼4 ms^-1^. Indeed, we find that our simulations stay near the starting structure and that our predicted energy transfer distributions agreed well with the major population seen experimentally but missed the minor population seen for constructs like lysozyme_44-150_ (Figure 3A).

**Figure 3:**
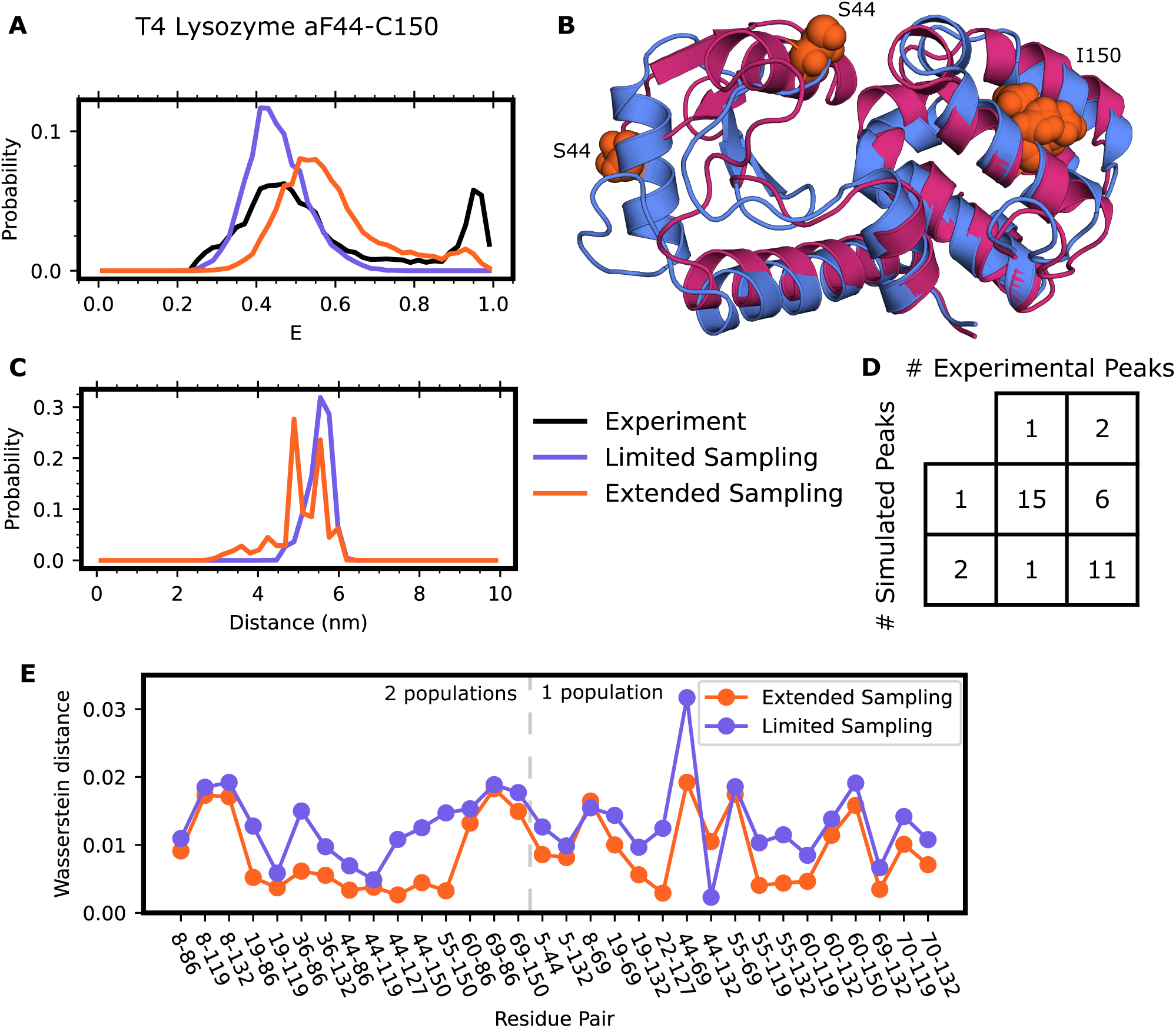
Discrepancies between smFRET and time averaging results enable discovery of a novel lysozyme fold. A) FRET efficiencies and (C) observed inter-dye distances for T4 Lysozyme labeled at position 44 and 150 with Alexa 488 and Alexa 647. Experimental traces are in black, calculations resulting from an MSM that only included crystal-like states in purple, and calculations resulting from an MSM including the alternate state in orange. Experimental donor only counts (E < 0.25) have been removed for ease of comparison. B) Example conformations of the crystal-like state of lysozyme (blue), or the alternate state (red). Residues 44 and 150 labeled for clarity. D) Qualitative comparison of smFRET distributions from experiment and simulation results including the alternate pose of lysozyme. E) Wasserstein distance between experimental and limited sampling dataset (purple) or extended sampling dataset (orange) for all labeled pairs.

To provide a structural explanation for the minor population seen experimentally, we employed a combination of metadynamics and MSMs. First, we used metadynamics simulations to find structures that are consistent with the high FRET of the minor population. In metadynamics, one adds an external biasing force to drive dynamics along a pre-selected collective variable. In this case, we pushed the system along the distance between residues 44 and 150 to see if we could find structures where they come close together, as this would result in a high FRET efficiency. These simulations revealed that the β domain can undergo minor unfolding which enables a swiveling motion to bring residue 44 much closer to residue 150 of the α domain (Figure 3B). To test if this alternative structural state is metastable, we selected four conformations where residues 44 and 150 are near one another and ran >500 ns conventional molecular dynamics simulations of each of them. All the simulations stayed near the starting point, confirming that the alternative structure state we discovered in metadynamics is a metastable free energy state. To determine the relative probability of this alternative state and those observed in our original simulations, we sought to build an MSM that captured transitions between the crystal-like states and the new alternative state. We ran goal-oriented adaptive sampling simulations using the Fluctuation Amplification of Specific Traits (FAST) algorithm^68^ to promote transitions from the crystallographic state to the alternative state and vice versa. FAST works by iteratively running a batch of simulations, building an MSM, and choosing states from the MSM as starting points for new simulations in a manner that balances between exploring further around states with a specified geometric property (called exploitation) and broad exploration of conformational space. In this case, we started one batch of FAST simulations from the crystallographic structure and set the exploitation term to favor states with a low RMSD to the novel fold we discovered and a second with the targets reversed. Both sets of simulations captured transitions between the two folds, providing a basis for building an MSM that captures the relative probabilities of both folds of lysozyme.

After building a new MSM that incorporates this data, we find that the computationally calculated energy transfer distribution now includes a minor state in agreement with experiments with multiple dye positions (Figure 3). Specifically, including the novel fold of lysozyme greatly improves agreement between our time-averaged results and the experimental traces for labeling pair 44-150 (Figure 3A,B). As a further test of our model, we then calculated energy transfer distributions for the remaining 32 FRET probe positions. Agreement between our model and these experiments would be strong support for our model, given that none of these experiments influenced our simulation strategy. In support of the alternative fold we predicted, we see the addition of minor peaks to 11 of the 17 FRET probe positions that were sensitive to this minor population, and only one additional peak in probe positions not reporting on the minor population (Figure 3D, S3). Inclusion of the alternate state greatly reduces the Wasserstein distance between the experimental and predicted histograms for all states except 2: 8-69 and 44-132.

### Results for IDPs are more sensitive to force field choice than globular proteins

We hypothesized that accounting for time-averaging in our Aβ40 prediction was insufficient to give strong agreement with experiment because the force field preferred overly compact states of Aβ40. Historically, force fields which govern the underlying physics of the simulation, have yielded large differences between simulations and the corresponding experimental data for IDPs ^63,69,70^. While amber99sb notably performs better with IDPs than others, force fields that were parameterized for folded proteins tend to lead to over compaction of IDPs, largely due to an imbalance between protein-protein and protein-water interactions. This systematic compaction of Aβ40 would skew the observed FRET values towards higher efficiencies, exactly as we have observed (Figure 2B).

To test whether discrepancies in experimental agreement are due to force field errors, we predicted Aβ40 smFRET using simulations conducted with a suite of nine different force fields/water models. Seven of these datasets were taken from a previous study that examined how well 30 μs simulations with each force field recapitulated NMR measurements. The seven force field/water combinations are: amber99SB*-ILDN with TIP3P, C22* with TIP3P-CHARMM, C36m with TIP3P-CHARMM, a03ws with TIP4P, a99SB with TIP4P-Ew with Head-Gordon vdW and dihedral modifications (a99sb-ucb), a99SB-ILDN with TIP4P-D, and a99SB-disp with a modified TIP4P water. In addition to these datasets, we also ran our own simulations using amber03^71^ and a99sb-ws, both with TIP3P water. For each of our simulations, we ran 250 ns long simulations in triplicate starting from the 10 most distinct Aβ40 structures captured in the previous simulations.

We find that force field choice substantially affects the quality of smFRET prediction for IDPs and that there is still room for improvement. For each of the above force field-water combinations, we generated MSMs and calculated the expected smFRET. All force field-water combinations result in Aβ40 distributions that are more collapsed than the experimental distribution (Figure 4A). We note that in our datasets, we explicitly started simulations from expanded states of Aβ40. However, these states exhibit a rapid compaction event which is not reversed during the simulation, consistent with previous findings that most force fields are biased towards more compact IDP structures than are experimentally observed ^42,69,72^. Of the force field-water combinations, a99SB-ILDN-TIP4PD and a99sb-ucb showed the strongest agreement with experiment.

**Figure 4:**
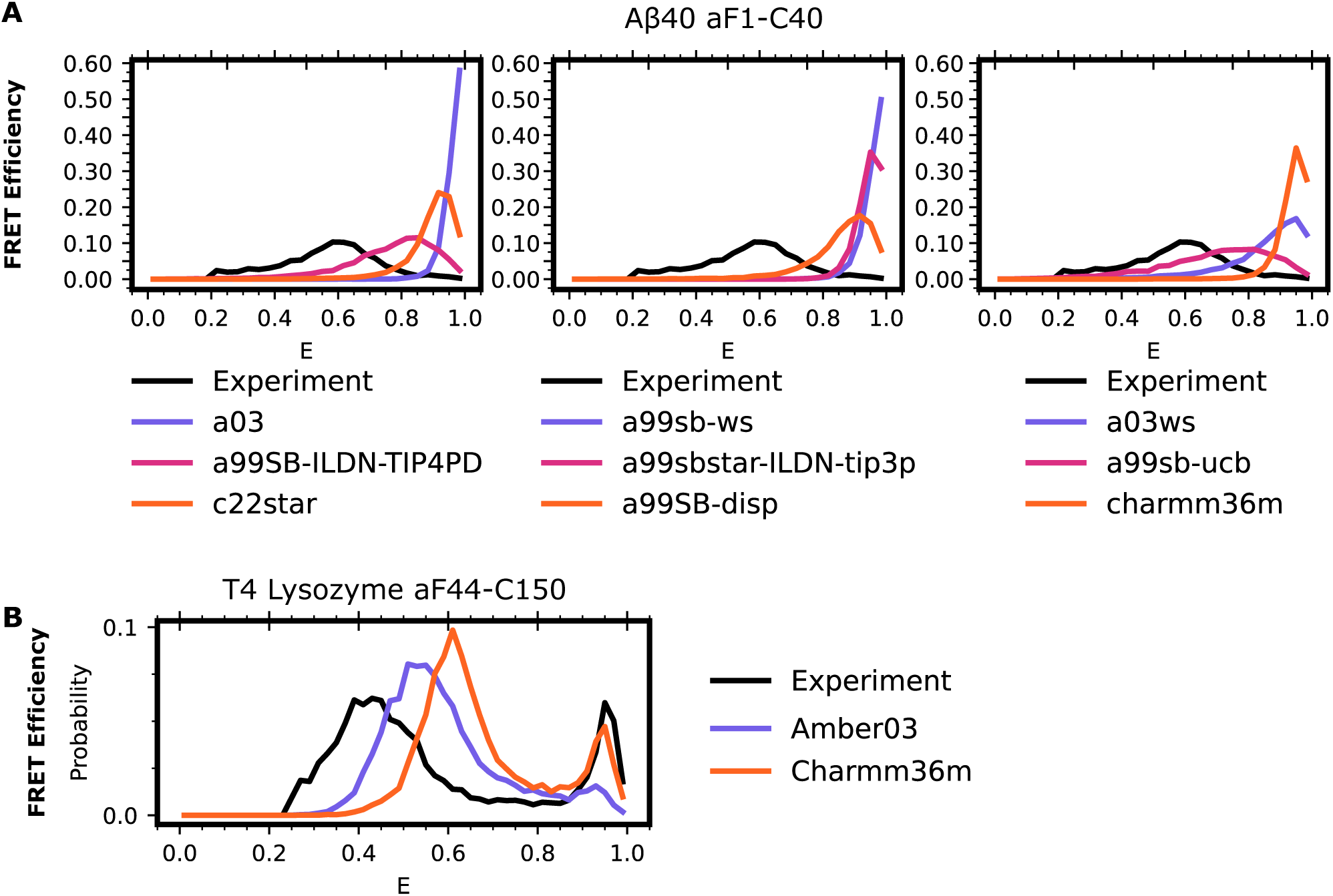
Force field choice has a significant impact on the level of agreement between simulations and experiments on IDPs even when accounting for time averaging. A) FRET efficiencies for Aβ40 labeled at positions 1 and 40 with Alexa 488 and Alexa 647 in different force fields. In each trace, the black line represents the experimental smFRET result, individual force field and water combinations indicated in the legend below the column. B) FRET efficiency distributions for T4-lysozyme residues 44 and 150 in amber03 with TIP3P water (purple) or charmm36m with TIP3P water (orange) or the experimental result (black). Experimental donor only counts (E < 0.25) have been removed for ease of comparison.

While there is large variation in force field performance for IDPs, we find less variation between force fields for lysozyme. We performed another 5 independent, 5μs long replicate simulations of lysozyme in tip3p water using charmm36m as a force field. As with our amber03 simulations, we fail to uncover the third, minor population, of lysozyme (Figure S4). Using the previously discovered minor state of lysozyme as both a starting and target structure, we again performed goal-oriented sampling to promote transitions between the crystal-like poses of lysozyme and the minor state in the charmm36m force field. Both sets of simulations again capture transitions between the two states, enabling us to construct an MSM. Though there are slight differences in energy transfer efficiency between our datasets constructed using amber03 and charmm36m, both produce good agreement between the experimental energy transfer distributions and those predicted from our MSMs (Figure 4B).

## Discussion/Conclusion

Here, we have explored how conformational averaging during smFRET measurements impacts the observed distribution of FRET efficiencies. Our results show that accounting for time-averaging across protein and dye dynamics improves the agreement between simulations and experiments for three proteins across the ordered spectrum (Figure 1, 2). These results agree with an existing experimental understanding that single molecule energy transfer distributions report an average of all protein motion during the measurement window. Our work adds to these prior experimental efforts by both identifying which states are being averaged together, while also providing an atomistic view of the protein conformations. While prior computational efforts have often focused on accounting for protein dynamics, improving protein and dye force field accuracy, or accounting for dye dynamics in FRET predictions, often these efforts either require additional simulations for every labeled dye position, or are unable to account for the effect of dye dynamics without *a priori* knowledge of the dynamical nature of the dyes. Our approach is unique in that it leverages MSMs to account for both protein and dye dynamics without the need for additional, computationally expensive, simulations. This removes additional modelling choices, such as choosing a Förster radius or timescale to average dye motions over, from the calculation.

Historically it has been difficult to determine why simulations and experiments have failed to agree. While simulations can have systematic errors due to parameterization or incomplete conformational sampling, experimental limitations and artifacts may also lead to disagreements. Here, we highlight examples where predicted and experimentally obtained energy transfer efficiency measurements appeared to disagree until we properly accounted for details of the experiment like time-averaging. Accounting for these experimental details in our modeling approach did not provide a structural rationale for a previously observed minor population of T4 lysozyme. However, we were able to use this persistent discrepancy to guide additional simulations to find structures that are consistent with the energy transfer in the minor populations. This approach allowed us to propose a structural model for the previously unexplained minor population that is consistent with most of the experimental measurements (Figure 3). We also find that simulations of Aβ40 with force field and water combinations parameterized for folded proteins result in an overly compact ensemble compared to the experimentally determined ensemble (Figure 4). As expected, based on previous publications, force fields designed to improve performance on disordered proteins performed better in our tests. While there were still differences between amber03 and charmm36m force fields for our lysozyme simulations, the choice of force field was less impactful for lysozyme than Aβ40.

We expect our approach will enable combining simulations and experiments to understand the link between sequence, structure, and function in many settings. While smFRET experiments are extremely valuable, one cannot readily derive atomistic models of conformational distributions from this data alone. The approach we have outlined here enables robust calculation of energy transfer distributions from protein ensembles, providing a direct link between energy transfer distributions and atomic models. While our approach led to strong agreement between simulations and experiments, there are some exceptions where our datasets diverge. One explanation could be that we do not consider alternative mechanisms of donor energy emission – such as quenching via nearby residues such as tryptophan and tyrosine. Another explanation could be that the mutagenesis required for dye attachment during experiments, as well as the attachment of the dye itself, disrupt the conformational landscape of the protein in question. Indeed, in many of our apo-simulation models we observe states that are incompatible with dye labeling, such as when amino acid to-be-labeled becomes buried or interacts closely with other regions of the protein. Nonetheless, our method is implemented post-simulation, modeling additional dye positions is rapid and requires minimal additional computational cost. Accordingly, once one has a satisfactory simulation dataset, it is facile to use these tools to design novel probe pairs which report on identified motions of interest.

## Brief Methods

### Molecular dynamics simulations

All simulations were performed in explicit solvent at 300K.

Simulations of Apolipoprotein E4 were generated using OpenMM8.0^73^ and amber03^71^ with TIP3P water^74^ and a timestep of 4 fs. The dataset was generated using a diverse composition of starting structures of ApoE and totals 3.61 ms. Clustering was performed based on the pairwise distances between 15 selected residue pairs and the MSM was created using row normalization and a 2 ns lagtime.

Simulations of Aβ40 in the following force fields were obtained from prior work: a99SB*-ILDN with TIP3P, C22* with TIP3P-CHARMM, C36m with TIP3P-CHARMM, a03ws with TIP4P/2005 interactions, a99SB with TIP4P-Ew with the Head-Gordon vdW and dihedral modifications (a99SB-UCB), a99SB-ILDN with TIP4P-D, and a99SB-disp with a modified TIP4P-D water. For each combination, a total of ∼30 μs data were collected using Anton hardware. Simulations of Aβ40 in amber03 + TIP3P and amber99sb-ws + TIP3P were generated in this work using GROMACS and 10 diverse starting structures from the above Aβ40 dataset and running each simulation for 250 ns in triplicate using unique initial velocities for each (aggregate 7.5 μs). Clustering for both simulation datasets was performed using the distance between every 5^th^ residue as a feature, and the MSM was created using row normalization and a 5 ns (Anton datasets) or a 0.2 ns lag time (GROMACS).

Simulations of T4 lysozyme were performed in amber03 with TIP3P water. Initial unbiased simulations were started from PDB structure 5LZM and 5 replicates were performed for 5 μs each using differing initial velocities. Metadynamics simulations were performed using PLUMED and a biasing potential of 0.3 between residues 44 and 150 for a total of 250 ns. Unbiased simulations were started from 4 alternate states uncovered by metadynamics with 5 replicas using differing initial velocities, each for a total length of 1 μs. FAST adaptive sampling was performed from both alternate and crystal-like states to the opposing state to capture the transition pathways in forward and reverse using RMSD as a progress metric. MSMs were built based on cluster centers from initial unbiased simulations, or the entire dataset excluding the metadynamics runs (total 94.8 μs). Clustering was performed using backbone RMSD to a radius of 2.5Å and the MSM was built using row normalization and a lag time of 2 ns.

Simulations of dyes were performed in amber03ws using the modified amber dye parameters^28,39^ with TIP3P water and a timestep of 2 fs. A single run was performed for each dye for 500ns. Simulation frames were saved every 20 fs. A 5000 state MSM with a lag time of 2 ps was built for each dye using RMSD of heavy atoms as a clustering metric.

### Simulation of smFRET

Dye color determination was achieved by building a MSM for each dye of interest (see simulations, above). Briefly, the dye MSM is modeled onto each state in the protein MSM, removing any positions from the dye MSM that clash with the protein. Next, a random dye position is chosen for both the donor and acceptor dye. Probabilities of radiative decay, energy transfer, non-radiative decay, or remaining excited are calculated and a random outcome is chosen accordingly, similar to prior work ^40,41^. If the dye remains excited, dye positions are allowed to update along with transfer probabilities until the donor dye is no longer excited. In the case of point clouds, a conformational ensemble of dyes ^26^ was modeled onto the protein and all steric clashes were discarded. Photon colors were determined by choosing a random distance between the donor and acceptor dye emission centers and the Förster relationship (Equation 1).

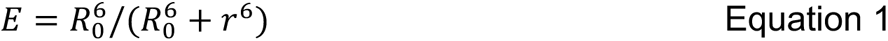

To determine which protein states to average, we recolor an experimental photon trace from Apolipoprotein E4 ^54^. We choose a random state from our protein MSM and build a synthetic trajectory to match the length of the experimental photon burst. We apply a time correction factor of 10,000 to slow the simulation timescale to match the experimental timescale. Each time an experimental photon is observed, we select the corresponding state in our trajectory and evaluate the photon identity as above. We determine the overall energy transfer efficiency as the ratio of acceptor photons to the total observed photons and repeat this process for all bursts (∼14,000), yielding the displayed distributions. The code for these calculations is available on github (https://github.com/bowman-lab/enspara).

### Experimental smFRET data

Data for Apolipoprotein E4 was obtained from Stuchell-Brereton *et al*^54^. Data for T4 lysozyme was obtained from Sanabria *et al*^55^. Data for Aβ40 was obtained from Meng *et al*^28^.

### Analysis/Software

Simulations generated during this manuscript were performed in GROMACS2020^75^ or OpenMM8.0^73^ as noted. Adaptive sampling was performed using FAST^68^, and metadynamics^76^ simulations were performed using PLUMED^77^ and GROMACS2020. Structure imaging was performed in PyMOL. Trajectory analysis was performed using MDtraj^78^. Clustering and MSMs were created using ENSPARA^79^. All graphs were generated using Matplotlib^80^.

## Acknowledgements

We thank the citizen scientists who contributed computational time and technical expertise for the simulation of ApoE via Folding@home. This work was supported by the National Institutes of Health through U19AG069701 (project 1: A.S, core B: G.R.B.), RF1AG067194 (G.R.B. and A.S.), NIGMS R35GM152085 (G.R.B.), and NSF MCB 2218156 (G.R.B.). J.J.M was funded by the NIH training grant T32AG05851804.

## Supplemental Methods

### Molecular dynamics simulations

#### Apolipoprotein E4

The simulation dataset for Apolipoprotein E4 was generated similar to Stuchell-Brereton *et al.* 2023^54^. Briefly, the NMR structure of an ApoE3-like protein (PDB: 2L7B)^81^ was used as a starting point. Mutations reverting this structure to the sequence of ApoE2 (112C, 158C), ApoE3 (112C,158R), ApoE4 (112R, 158R), and ApoE3-Christchurch (112C, 158R, 136S), and each mutation underwent 20 rounds of adaptive sampling using the FAST algorithm to explore the three distances pairs: R92 and S263, G182 and A241, and S223 and A291. All datasets were clustered into a shared model using backbone RMSD to a minimum difference of 3.5Å, yielding 18,182 centers. Each cluster center was solvated in a dodecahedron box with a 1.0 nm pad from the largest observed cluster center containing 0.1M sodium chloride. Each center was energy minimized and equilibrated by starting simulations at 20K and heating to 300K over a period of 2ns before a final NPT equilibration at 300K of 0.4ns. Each structure was launched on Folding @ home twice using different initial velocities and each trajectory reached 100ns, yielding an aggregate simulation time of 3.61 ms. All simulations were performed in the amber 03 force field with TIP3P water, hydrogen mass partitioning at 300K and a timestep of 4 fs. FAST simulations were performed using GROMACS and Folding @ home simulations were performed using OpenMM. Simulations were clustered using distance based clustering using the 5 FRET probe positions and 10 additional residue pairs as features, to generate a coarse model containing 8000 cluster centers. A Markov State Model was generated using a 2 ns lag time and ENSPARA’s row normalization builder.

#### Aβ40

Simulations of Aβ40 were acquired from Robustelli *et al.* Briefly, an extended conformation of Aβ40 was simulated in the following force fields: a99SB*-ILDN with TIP3P, C22* with TIP3P-CHARMM, C36m with TIP3P-CHARMM, a03ws with TIP4P/2005 interactions, a99SB with TIP4P-Ew with the Head-Gordon vdW and dihedral modifications (a99SB-UCB), a99SB-ILDN with TIP4P-D, and a99SB-disp with a modified TIP4P-D water. Simulations were run at 300K in NPT ensemble on Anton hardware with a 2.5-fs time step for a total of ∼30 μs.

For simulations generated during this manuscript, we used either amber03 or amber99sb-ws force fields with TIP3P water. Simulations were started from the top 10 divergent structures of Aβ40 found in the Robustelli *et al.* simulations. Each structure was solvated in a cubic box with box lengths of 12.307 nm which was determined by solvating the fully unfolded Aβ40 with a 1 nm pad, 0.1M sodium chloride, and virtual sites for hydrogens. Each structure was energy minimized and allowed to equilibrate for 1 ns before starting production runs. Three 250ns long replica simulations with independent velocities were started from each pose, for a total of 7.5 μs of aggregate simulation time using a 4 fs timestep.

Clustering was performed using the distance between every 5^th^ residue as an input feature to generate 250 unique cluster centers. MSMs were generated using a lagtime of 5 ns (Anton datasets) or 2 ns.

#### T4 Lysozyme

Simulations of T4 Lysozyme were initialized using PDB structure 5LZM, solvated in a cubic box with edges extending 1.8 nm beyond the edge of the protein with TIP3P water and 0.1M sodium chloride. Virtual sites were included for hydrogens. Structures were energy minimized for 1000 steps and equilibrated for 1 ns prior to production runs. For initial unbiased simulations, 5 replica production runs were performed for 5 μs each with each run having differing initial velocities.

Metadynamics simulations were performed using PLUMED with the metad restraint using a pace of 500, gaussian height of 0.3, and gaussian widths of 0.05 for a total of 250 ns. Biases were placed on the distance between residue 44 and 150 using both CA-CA distance the terminal side-chain atoms. 4 divergent structures were taken from the minimal 44-150 distances observed in the metadynamics simulation, resolvated in a cubic box with a 1.8 nm pad of TIP3P water and 0.1M sodium chloride, and re-energy minimized and re-equilibrated. Each pose was run with 5 replicates with differing initial velocities for 1 μs per each replica.

Adaptive sampling simulations exploring the transition between 5LZM and the alternate state of lysozyme identified by metadynamics were performed using FAST. Briefly, 10 40 ns long simulations were started from either 5LZM or the alternate pose of lysozyme. These simulations were clustered, a MSM was built, and 10 states with a minimal backbone RMSD to the target state were chosen to restart 40 ns simulations from. We iterated between clustering and simulation until states were identified with a backbone RMSD of <2Å. All simulations were performed at 300K with GROMACS 2020.

Coarse grained models were built on the initial unbiased simulations from 5LZM or a combination of the initial unbiased simulations from 5LZM, the 4 differing alternate states, and the FAST simulations observing the transition between the alternate state of lysozyme and 5LZM. For both models, clustering was performed based on RMSD of the backbone to either 500 centers, or a final RMSD of 2.5 Å, whichever was greater. Markov State Models were generated using ENSPARA’s normalize method with a lagtime of 2 ns.

#### Post-simulation modeling of smFRET

Direct simulation of dye emission events was achieved by building a MSM for both donor and acceptor dyes. We model each dye conformation from the MSM onto each labeling position in the protein MSM, discarding any dye positions that resulted in steric clashes with the protein. Next, for each state in the protein MSM, we simulate dye emission events similar to previous methods ^40,41^. Briefly, we choose a random dye starting position for both the acceptor and donor dyes from the MSM based on their equilibrium probability. Next, we calculate the probability that the donor dye can undergo radiative decay (p_rad_, emit a donor photon), transfer energy to the acceptor (p_RET_, emit an acceptor photon), non-radiatively decay (p_nonrad_ no observed photon), or remain excited (p_remain_).

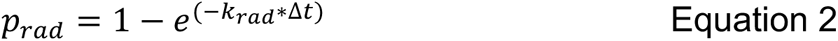

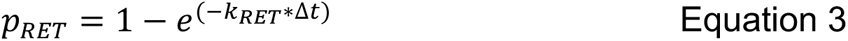

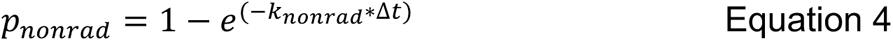

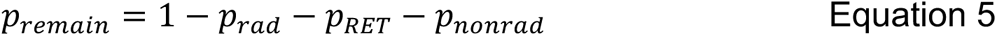

Where Δt is the timestep of the Monte Carlo which is the same as the dye MSM lagtime (2 ps), k_rad_ is the rate of radiative decay, k_RET_ is the rate of energy transfer, and k_nonrad_ is the rate of nonradiative decay, given by the following:

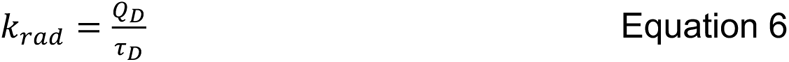

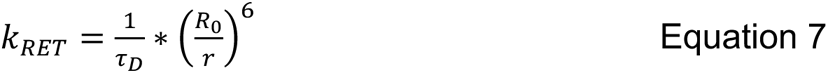

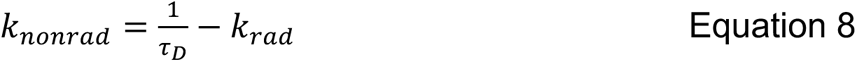

Where Q_D_ is the donor fluorescence yield in the absence acceptor, τ_8_ the donor lifetime in the absence acceptor, r the distance between the donor and acceptor emission centers, and R_0_ the Förster radius, given by the following:

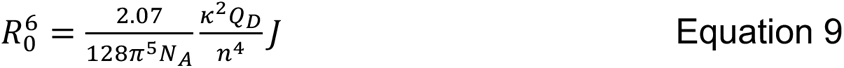

Where N_A_ is Avogadro’s number, n the refractive index of the solution, J the donor-acceptor spectral overlap integral, and κ^2^ the dipole orientation factor between the two dyes. In this equation, we hold all values constant except for κ^2^ which we calculate according to:

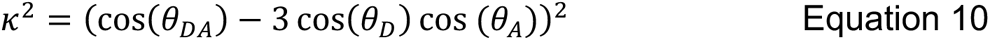

Where θ_DA_ is the angle between the donor dipole moment and the acceptor dipole moment, θ_D_ the angle between the donor dipole moment (*d̂*) and the vector connecting the donor emission center to the acceptor emission center (*r̂*), and θ_A_ the angle between the acceptor dipole moment (*â*) and *r̂*.

After we calculated the emission outcome probabilities, we choose a random outcome weighted by the respective probability of occurring. If the donor dye remains excited, we allow both dyes to update their positions based on the probability of transitioning states from the dye MSM, recalculate potential emission outcomes, and choose another dye outcome. We repeat this process until the donor dye is no longer excited, recording both the number of Monte Carlo steps required to reach the emission event (dye lifetime) as well as the outcome. To enable efficient computation, we pre-calculate the lifetimes and outcomes for each protein center, repeating each Monte Carlo simulation 1000 times (ApoE) or 5000 times (Lysozyme, Aβ40) to scale with the respective numbers of cluster centers that each protein MSM has.

For the dye point cloud method dye molecules of interest were modeled onto the protein at the appropriate labeling positions. We do this using a rotamer library approach based on prior work^26^. Briefly, dyes attached to the appropriate label and linker were simulated free in solution to determine all the potential dye configurations. All resulting simulation frames were aligned based on the backbone and the center of fluorescence emission from the dye was saved as a single point to generate a point cloud of all potential emission centers. Next, the point cloud is modeled onto the protein labeling position of interest and all points that would result in a steric clash are discarded. Finally, we generate a distance probability distribution which describes the distance between all potential configurations of the donor and acceptor dyes. We determine the photon color by choosing a random donor-acceptor distance, assessing the probability of transfer based on the Förster relationship (Equation 1), and choose whether the donor was a photon or acceptor based on the established probability. We specify an R_0_ of 5.6 nm for Alexa488 and Alexa594.

To account for protein conformational sampling during the measurement window, we recolor an experimental photon time course from experiments performed on ApoE. Using our MSM, we generate a synthetic trajectory that matches the length of the experimental photon burst. The trajectory starts from a random state in the MSM based on the equilibrium probability of that state, and the synthetic trajectory is built based on the probability of that state transitioning to any other state in the MSM. The length of the trajectory is determined based on the length of the experimental photon burst and rescaled to account for simulations being faster than experiment. In our calculations, we use a time-factor of 10,000. Each time an experimental photon is recorded, we note the corresponding frame in the synthetic trajectory and decide whether the photon was a donor or acceptor photon based on the schema outlined above (dye emission simulation or point cloud approximation). We repeat this for each observed photon in the experimental burst and return a total FRET efficiency for the burst as the ratio of observed acceptor photons and total observed photons. This entire process is repeated, generating new synthetic trajectories from new starting states, for each observed molecule in the ApoE experiment resulting in >14,000 observations.

Code used to run dye modeling and smFRET calculations are available on github: https://github.com/bowman-lab/enspara. MSMs of proteins and dyes, as well as example code for running smFRET calculations and generating dye MSMs is available on OSF: https://osf.io/82xtd/?view_only=b7f354e86eb144a69d9d047b42e21a9f.

#### Analysis/Software

Simulations generated during this manuscript were performed in GROMACS or OpenMM as noted. Structure viewing was performed in PyMOL. Trajectory analysis was performed using MDtraj. Clustering and MSMs were created using ENSPARA. All graphs were generated using Matplotlib.

### Supplementary Figures

**Figure S1:**
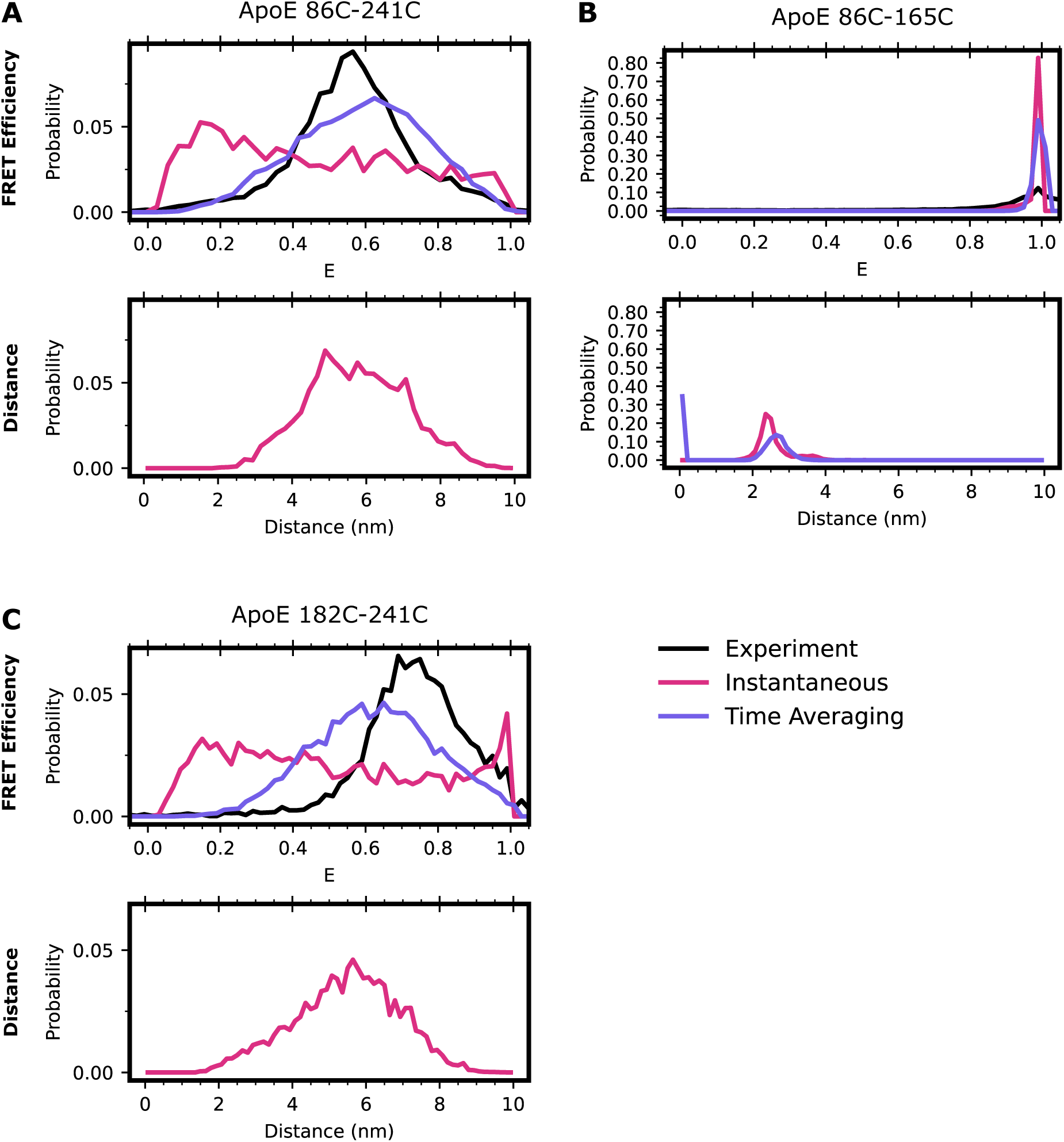
Accounting for time averaging significantly alters the apparent structural distribution from our model and increases agreement with experiments. Top, smFRET histograms for experimental (black), instantaneous simulation (red), or time averaged simulation (purple), bottom the inter-dye distances for apolipoprotein E. Labeled positions are A) 86-241, B) 86-165, or C) 182-241. In all cases, labeling is performed with Alexa 488 and Alexa 594 using maleimide chemistry.

**Figure S2:**
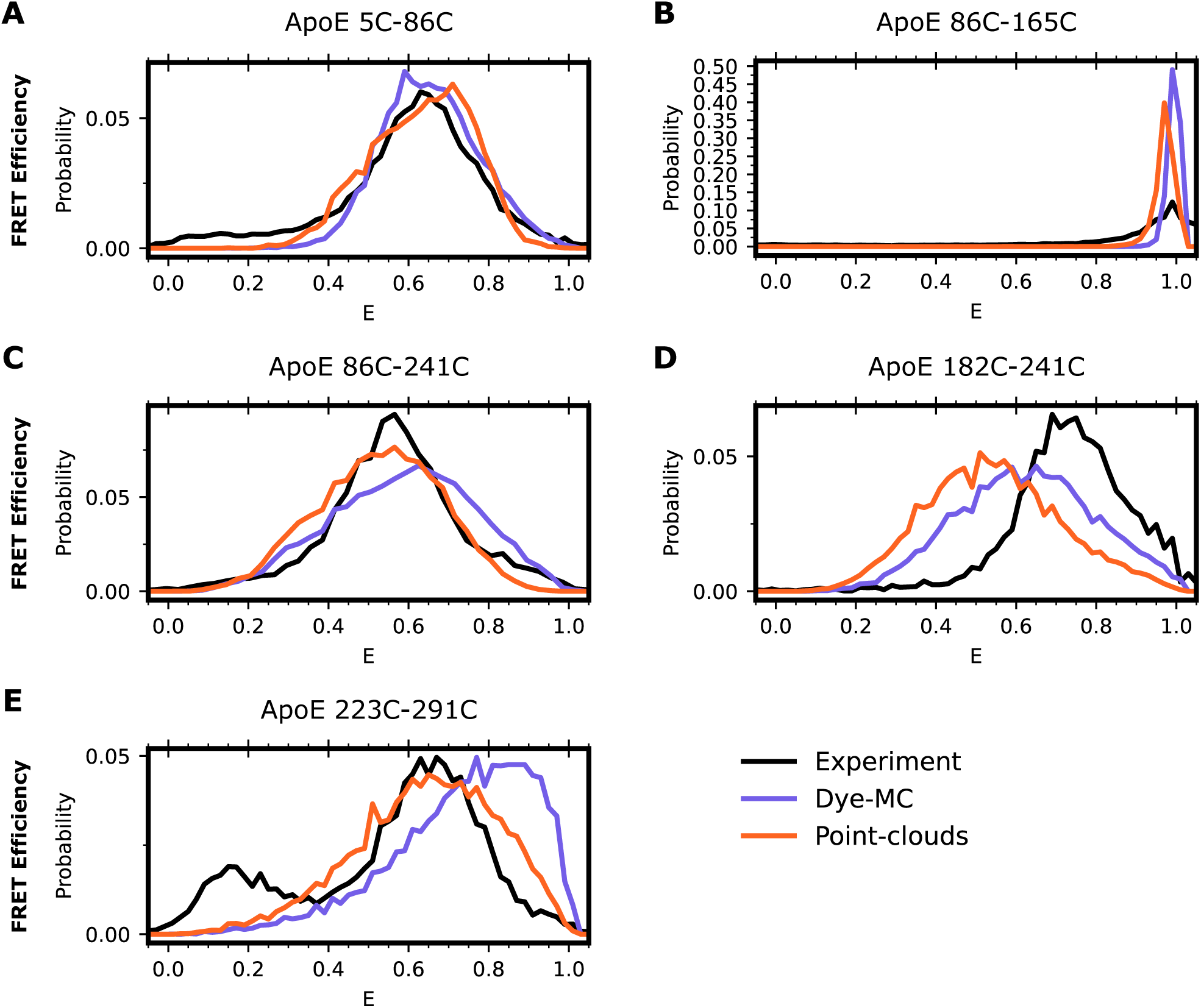
Treating dyes as point clouds yields comparable results to accounting for dye dynamics. FRET efficiencies for apolipoprotein E labeled with Alexafluor 488 and Alexafluor 594 at positions A) 5-86, B) 86-165, C) 86-241, D) 182-241, and E) 223-291. The black trace is the experimental distribution, in purple is accounting for time averaging while accounting for dye-dynamics (Dye-MC), and in orange is treating dyes as a point cloud with no dynamics, using a constant R0 of 5.6.

**Figure S3:**
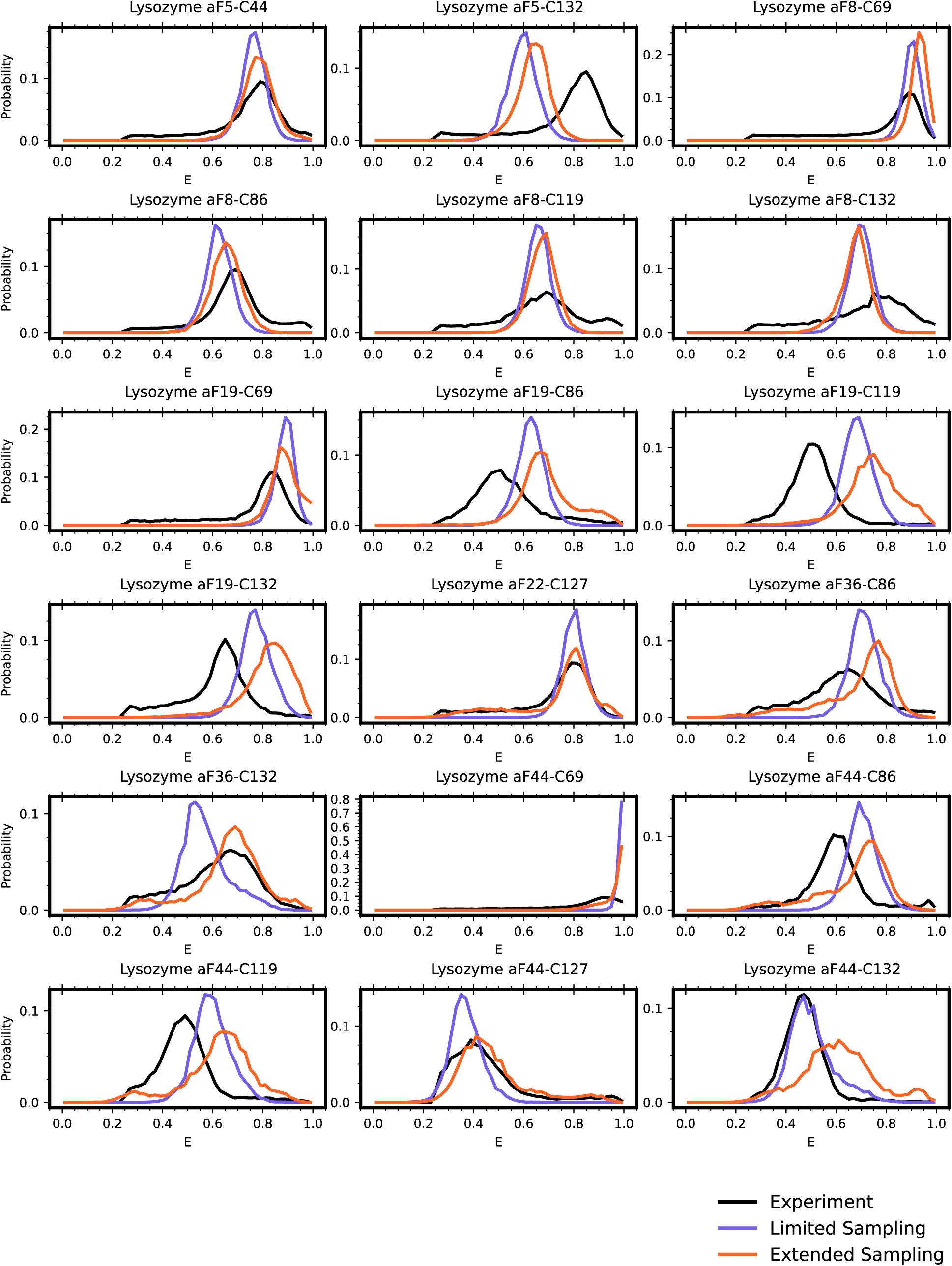
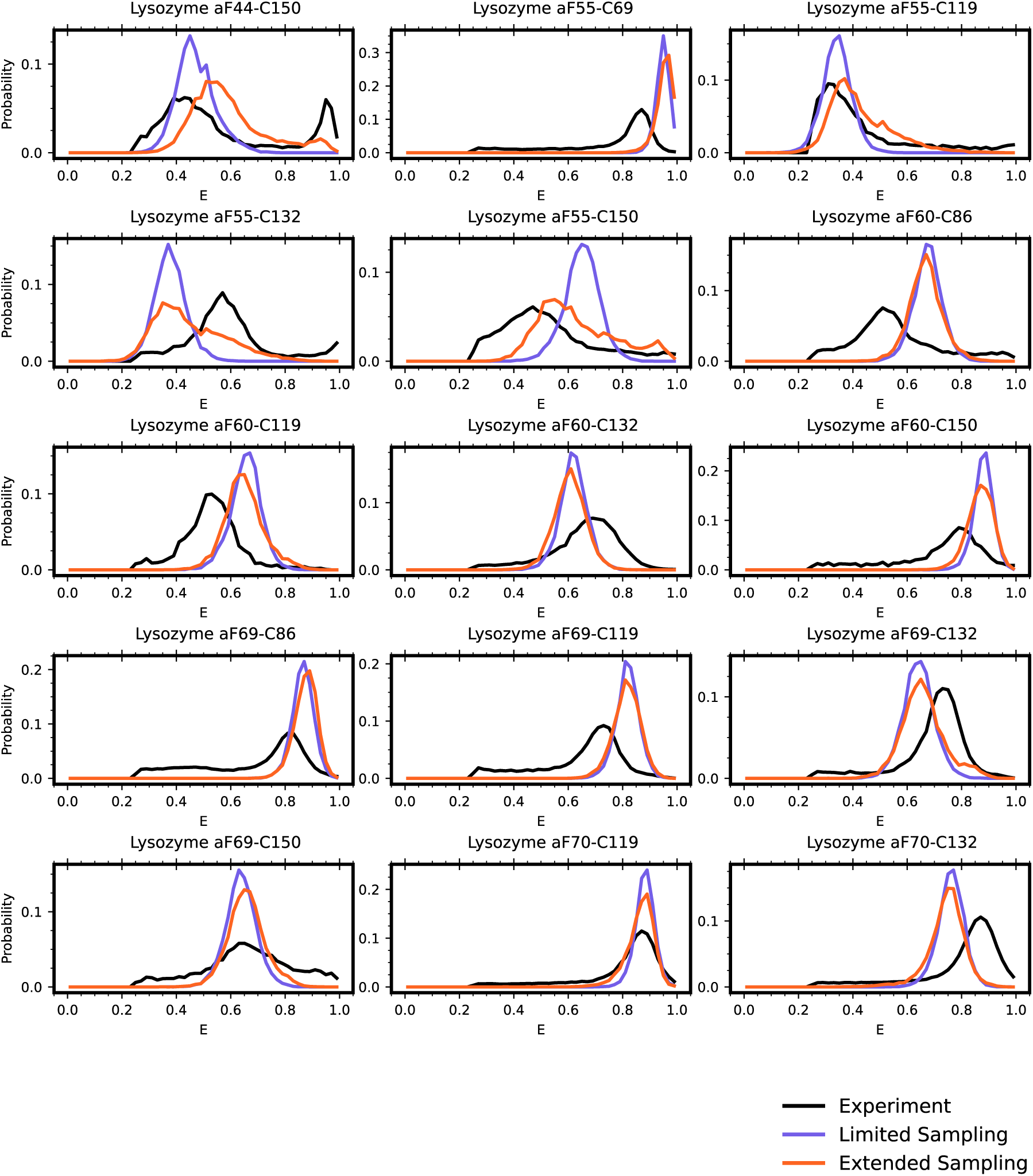
Extended sampling of lysozyme yields improved agreement with experiment. FRET efficiency distributions for various lysozyme probe positions. In black is the experimental trace, donor only counts (E < 0.25) have been removed for comparison purposes as simulated FRET has total labeling. In purple is the distribution from our initial simulation runs which only sample crystal-like poses. In orange is a model of Lysozyme which includes the novel state. Simulations run in amber03 force field using TIP3P water. All calculated FRET was performed while accounting for both dye and protein dynamics.

**Figure S4:**
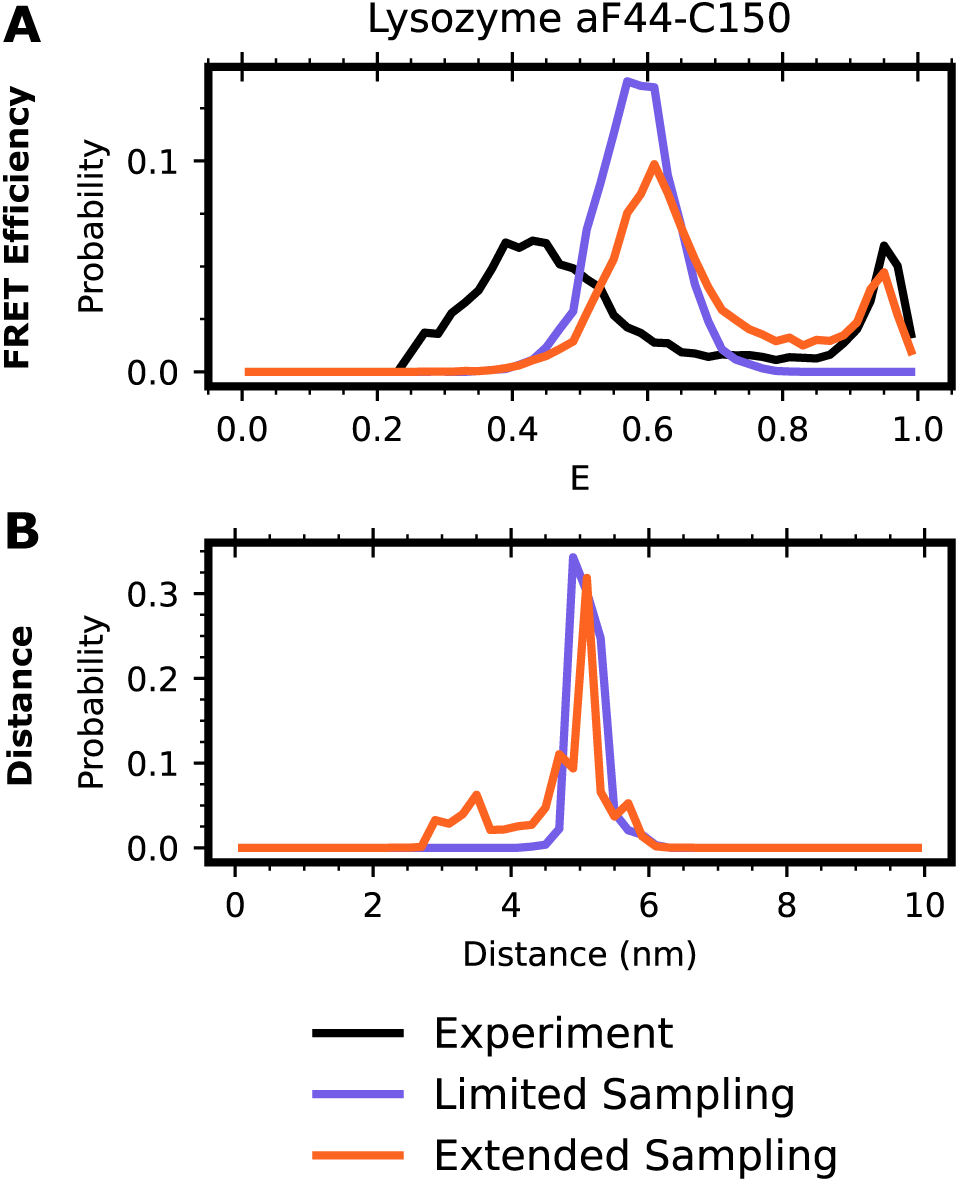
Limited sampling of lysozyme using charmm36m fails to recapitulate the third state of Lysozyme. FRET efficiency distributions for lysozyme 44-150. The black trace is the experimental distribution with donor only counts (E < 0.25) removed for clarity. in red is the equilibrium distance distribution from simulation accounting for added dye-distances, and in purple is the effect of time averaging on the red trace. Simulations run in charmm36m with TIP3P water.

